# Multidisciplinary Interrogation of a Crucial Protein Interface in the Type II Secretion System

**DOI:** 10.1101/2020.12.11.420943

**Authors:** Cristian A. Escobar, Badreddine Douzi, Geneviève Ball, Brice Barbat, Sebastien Alphonse, Loïc Quinton, Romé Voulhoux, Katrina T. Forest

## Abstract

The type IV filament superfamily comprises widespread membrane-associated polymers in prokaryotes. The Type II secretion system (T2SS), a significant virulence pathway in many pathogens, belongs to this superfamily. A knowledge gap in the understanding of the T2SS is the molecular role of a small ‘pseudopilin’ protein. Using multiple biophysical techniques, we have deciphered how this missing component of the Xcp T2SS architecture is structurally integrated, and thereby also unlocked its function. We demonstrate that the low abundance XcpH is the adapter that bridges a trimeric initiating tip complex XcpIJK with a periplasmic filament of XcpG subunits. Our model reveals that each pseudopilin protein caps an XcpG protofilament in an overall pseudopilus compatible with the dimensions of the periplasm and the outer membrane-spanning secretin through which substrates of the T2SS pass. Unexpectedly, to fulfill its adapter function, the XcpH N-terminal helix must be unwound, a property shared with the XcpG subunits. We provide the first complete structural model of a type IV filament, a result immediately transferable to understanding of other T2SS and the type IV pili.

## INTRODUCTION

Bacteria have sophisticated secretory nanomachines that evolved to deliver exoproteins to the bacterial cell surface, into the surrounding medium or directly into host cells. Among these, the type 2 secretion system (T2SS) is a trans-envelope apparatus specialized for secretion of folded proteins from Gram negative bacteria^1^. In many cases, these secreted substrates are virulence factors: examples include the cholera toxin of *Vibrio cholera*, the exotoxin A of *Pseudomonas aeruginosa*, and the heat labile toxin of enterotoxigenic *Escherichia coli*^2^. The conformational constraint of transporting folded substrates is shared by the Type VII, or ESX, system^3^ and the Type IX secretion system^4^, but is in clear contrast to the linear export of unfolded polypeptides by the T3SS^5^ or the T4SS^6^. The substrates of the T2SS are not known to share any sequence or 3D structural motifs^7^, and thus an open question is how substrates are individually recognized and ushered out. Dynamic and changing interactions, potentially involving nucleation of structure in intrinsically disordered regions, must be at the heart of this selection process^8^.

The T2SS accomplishes its task with a distinctive mode of transport involving a pilus-like structure (the pseudopilus) in the periplasmic space, at the interface between an inner membrane assembly platform and a large outer membrane pore, the secretin^9^. Structural, functional and evolutionary commonalities to the extracellular Type IV pili critically inform the T2SS field and *vice versa*^10^. Thanks to recent spectacular developments in cryo-electron microscopy, the 3D structures of T2SS secretins^11–14^ and the filament formed upon overexpression of the pseudopilin subunit PulG^15^ have been solved, and reveal important structural features. While secretins form a tightly gated giant double *β*-barreled pore of 80 Å diameter in the outer membrane, PulG adopts a right-handed homopolymeric helix with a diameter of 70 Å, compatible with observations that a filament extends through the secretin when the pseudopilin is overexpressed^16,17^. In this pseudopilus, inter-subunit contacts occur along hydrophobic N-terminal *α* helices of the pseudopilin, with a special role for the negatively charged glutamic acid at position 5 in both recruiting and stabilizing interactions within the filament^18–21^. It has long been assumed that under native expression levels the pseudopilin forms a short and transient thread entirely within the periplasm during secretion^22^, a model which has been difficult to verify experimentally until recently, when a cryo-electron tomography reconstruction of the *Legionella pneumophil*a T2SS revealed pseudopilus density in ∼20% of the complexes^23^.

In addition to the major pseudopilin, additional low abundance (or *minor)* pseudopilins have been identified in the T2SS, based on being encoded within the same operon and sharing amino acid sequence with the pseudopilin, plus undergoing maturation by the same prepilin peptidase^24–26^. The four minor pseudopilins are generally designated as the H, I, J, and K proteins (variously prefixed with Gsp-, Pul-, Xcp-, depending on the particular T2SS, please see on line Materials and Methods for nomenclature notes). While none of these has been directly observed in a T2SS pseudopilus, the periplasmic domains of minor pseudopilins GspI, GspJ, and GspK have the ability to assemble into a ternary complex whose crystal structure has been solved. The four periplasmic domains of GspH, GspI, GspJ and GspK, form a quaternary complex^27,28^, but its structure is unknown. As a group, the minor pseudopilins are known to play a role in initiation and control of pseudopilus assembly and potentially retraction. The complex furthermore interacts with secretion substrates^32^ and with the periplasmic domains of the secretin^33^. Although there is strong evidence that the quaternary complex is found, at least transiently, at the tip of the pseudopilus, its structure has been elusive. Minor pseudopilin subunits could not be resolved in the *L. pneumophila* T2SS cryo electron tomograms^23^. In a recent T4P cryo electron tomography study, minor pilins were attributed to density blobs but no conclusions could be drawn about which protein belonged to which density nor about the molecular nature of their interactions^34^

While many reports have established the importance of the minor pseudopilins and pilins as a group for initiation and/or assembly of (pseudo)pili, fewer have analyzed their individual contributions to the secretion process. Nonetheless, specific roles have been assigned or proposed for all four minor pseudopilins: GspJ primes filament assembly through the recruitment of other minor pseudopilins^37^, GspI serves as a protein-protein interaction hub^27^, and GspK controls pseudopilus length and/or number^27,37^. GspH interacts with GspJ through its globular domain and with the structurally similar major pseudopilin via its N-terminal hydrophobic α-helices^18,27,39,40^; thus it is tempting to suggest GspH plays an adapter function between the tip and the body of the pseudopilus. However, the essentiality of GspH for secretion has been called into question because its absence can be overcome by overproduction of other minor pseudopilins^37^. There are no molecular data on how GspH is embedded in the quaternary complex.

Based on x-ray crystallographic structures of constructs representing periplasmic domains of GspH, GspI, GspJ, and GspK from several microbial species solved alone and/or in dimeric or trimeric subcomplexes^28,39–43^, each pseudopilin fold resembles major T2SS pseudopilins and Type IV pilus (T4P) major pilins characterized by an extended N-terminal *α*-helix, the first half of which (*α*1N) is not included in the soluble crystallography targets but is expected to be exposed to the membrane environment, and the second half of which (*α*1C) is partially buried in a globular α/β domain. Despite this common overall topology, it is important to bear in mind that the periplasmic domains of the minor pseudopilins interact to form the soluble quaternary complex in the absence of *α*1N, and since these minor subunits are at the tip of the pseudopilus they cannot depend on vertical interactions with major subunits for upward assembly interactions. Within this general framework, each minor pseudopilin has distinctive sequence and structural hallmarks. GspH is most similar in size, sequence and structure to the major pseudopilin. GspI is the smallest of the four. GspJ is larger, and displays a pronounced groove on its surface. Finally, GspK carries an additional structural domain of over 100 amino acids. GspH, GspI, and GspJ share the glutamate at position 5 with GspG, whereas GspK does not.

Based on two crystal structures of ternary complexes of minor pseudopilin periplasmic domains (3CI0 from the enterotoxigenic *E. coli* Gsp T2SS and 5VTM from the *P. aeruginosa* Xcp T2SS equivalent minor pilins XcpIJK^28,40^), we know the stabilizing interactions among the periplasmic domains (designated here with a subscript “p”) GspI_p_, GspJ_p_, and GspK_p_ include buried salt bridges between *α*1C residues for all three pairwise interactions: XcpI_p_ Asp51 to XcpK_p_ Arg45, XcpJ_p_ Glu58 to XcpK_p_ Arg45, and XcpI_p_ Asp51 to XcpJ_p_ Glu58. The interaction is also stabilized by salt bridges involving amino acids of XcpI_p_ *α*1 and XcpJ_p_ *β*11, including XcpI_p_ Asp51 and XcpK_p_ Arg195. Contacts at the bottom of the bundle of helices further lock the ternary complex in place. Double involvement of XcpI_D51_ in XcpI_p_:J_p_ & XcpI_p_:XcpK_p_ interfaces is evidence to bolster the original finding that XcpI_p_ supports the linear organization of XcpJ_p_:I_p_:K_p_ interactions in the complex^27^. Somewhat perplexingly, the largest buried surface area between any two of the soluble proteins in both ternary complexes is between XcpJ_p_ (GspJ_p_) and XcpK_p_ (GspK_p_), with a total buried surface area of ∼3600 Å^2^ in 5VTM. And yet, these two subunits do not interact with each other in solution in the absence of XcpI_p_^27^.

Several lines of evidence support downward addition of XcpH to the base of the XcpJ:I:K complex through an interaction between the globular domains of XcpH and XcpJ. XcpH_p_ completes the quaternary complex by interacting with XcpJ_p_^27^. Additionally, *α*1 of XcpH is essential in this interaction and XcpH_p_ can be positioned most convincingly in a small angle x-ray scattering envelope of the quaternary complex below XcpJ_p_^28^. The structural compatibility of the ternary tip complex GspI_p_J_p_K_p_ with downward addition of GspH below GspJ was proposed by Korotkov *et al*. in agreement with their previous suggestion that GspH could be positioned at the top of a GspG fiber but not at the bottom^39,40^. The lack of Glu5 in GspK can also be taken as evidence that GspK does not require helix:helix packing stabilization via a salt bridge with a higher subunit in the filament and thus is located at the top of the pseudopilus^31^.

In this study we set out to validate the biological importance of XcpH and to decipher the molecular details of the XcpH:I:J:K interaction. Because all available data to date suggest an unstable, dynamic and/or small interaction interface with XcpH, we turned to a collection of compatible techniques suited to interrogation of such interactions. We show that XcpH is indeed critical for efficient Type II secretion in *P. aeruginosa*. We demonstrate it interacts specifically with the *α*1 helix of XcpJ, and we narrow the interaction residues to a well-defined interface that is highly conserved among T2SSs. Our experimentally determined constraints allow us to synthesize a molecular model of the XcpHIJK quaternary structure, and to place this in the context of the complete pseudopilus filament, with biological implications for the role of the GspH/XcpH low abundance pseudopilin. Moreover, the molecular organization involving an inner membrane platform, priming by minor pilins, and a helical subunit assembly that is remarkably well conserved both evolutionarily and structurally in other type IV filamentous nanomachines^10,44^ including the canonical type IV pilus (T4P)^19,31,38^, competence pilus^45,46^, and to some extent the archaellum^47–49^. Thus our work has wide-ranging implications for understanding assembly of several fundamental microbial organelles.

## RESULTS

### The four minor pilins are essential for secretion

In our quest to understand the structure-function relationship of the minor pseudopilin complex, we endeavored to systematically assess the independent requirement of each of the four minor pseudopilins of the *P. aeruginosa* Xcp T2SS during the secretion process. Individual in-frame deletion mutants were constructed in the PA01 reference strain for *xcpH, xcpI, xcpJ* and *xcpK* (see online Materials and Methods for nomenclature). Each mutant was assessed for its ability to secrete the major Xcp T2SS effector LasB (**Figure 1a**) and for the ability of secreted LasB to degrade casein on milk plates (**Figure 1b**). All four individual deletion strains were similar in their inability to secrete LasB and lacked protease activity, like the negative control strain in which the entire T2SS operon is missing, thus demonstrating the absolute requirement of each minor pseudopilin in the Xcp T2SS secretion process. This finding is supported by complementation of each secretion deficient phenotype by introduction of the corresponding wild type gene on a plasmid. Our data are apparently in conflict with a 2018 publication suggesting that XcpH and XcpK are not required for T2S in *P. aeruginosa*^28^. Our finding that XcpH is indispensable for secretion prompted us to further explore the 3D integration of XcpH into the XcpHIJK quaternary complex.

**Fig 1.**
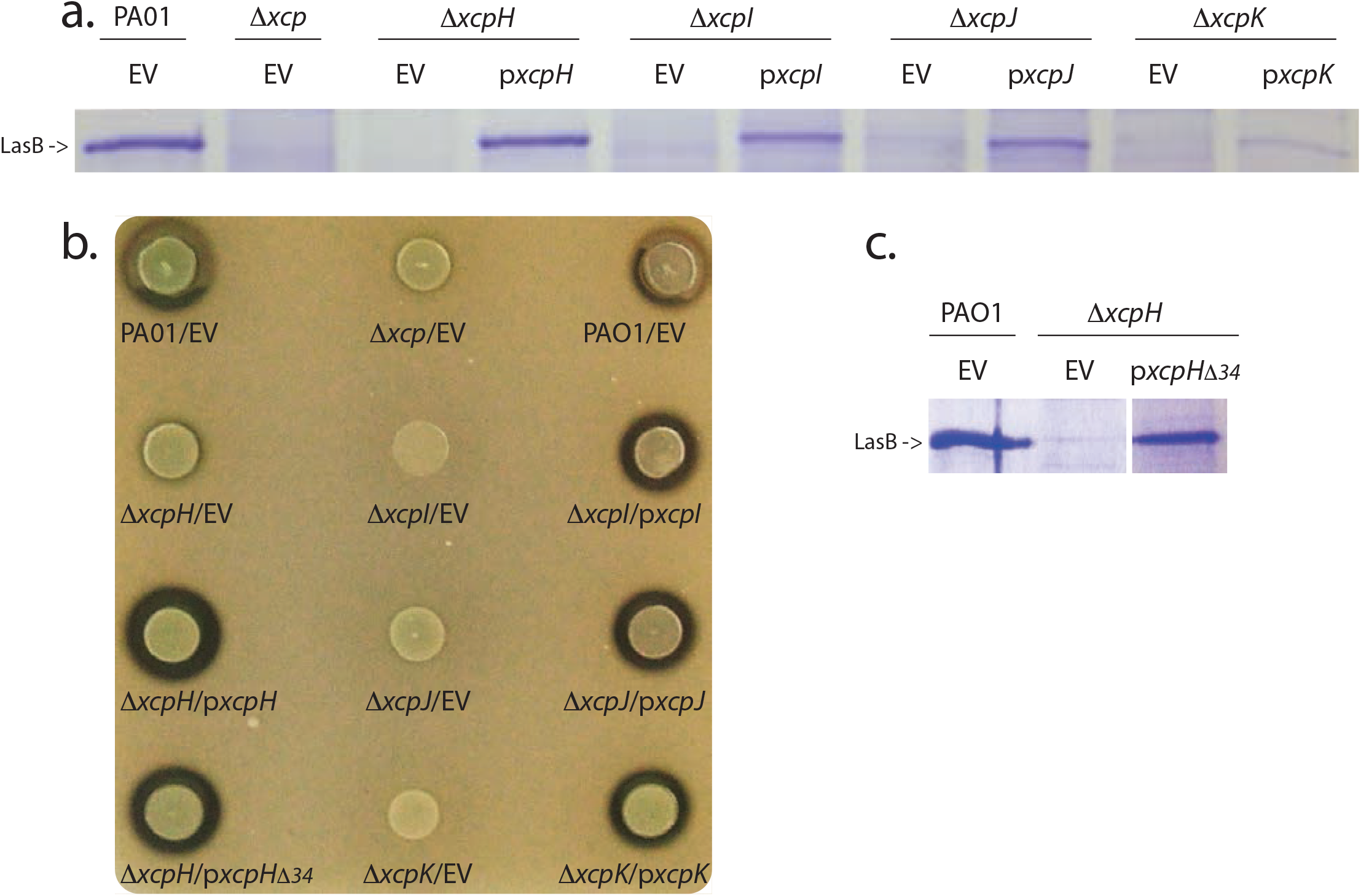
Minor pseudopilins are required for Type II Secretion. a. Secretion of the principal Xcp T2SS-dependent exoprotein (LasB) assayed by Coomassie blue stained SDS-PAGE of equal OD equivalent loading of culture supernatants of the wild type PAO1, Del xcp (DZQ40), Del xcpH, Del xcpI, Del xcpJ and Del xcpK strains harboring empty vector (EV), or a complementing plasmid encoding XcpH (pxcpH), XcpI (pxcpI), XcpJ (pxcpJ) or XcpK (pxcpK). b. LasB protease activity on a skim milk plate is indicated by a halo around a colony, corresponding to casein degradation by secreted LasB. Colonies representing WT (PAO1), Del xcp, or complemented Del xcp strains are labeled accordingly. Note Del xcpH is complemented by both pxcpH and by pJN105 expressing the XcpH deletion of the beta3-beta4 loop, pxcpHdel34. c. Secretion assay, as in panel a. showing secretion of LasB by Del xcpH/ pxcpHdel34.

### Chemical shift perturbation studies of labeled XcpH_p_ in the presence of XcpJ_p_

#### XcpH_**p**_ solution NMR assignments

To discover sites of interaction between XcpH and XcpJ, we performed solution NMR experiments. We initially isotopically labeled the ∼16 kDa XcpH_p_, a feasible NMR target, with ^15^N and ^13^C, and carried out amino acid assignments. Regions heavily crowded in the HSQC spectrum were excluded, leading to 85% coverage of the XcpH_p_ amino acid sequence (**Figure 2a**). The high signal dispersion of the HSQC spectrum indicates an overall well-folded protein, while the area in the spectrum with high resonance overlap and strong signal (around 8.2 ppm in the ^1^H dimension and 121 ppm in the ^15^N dimension) indicates a highly dynamic region of the protein.

**Fig 2.**
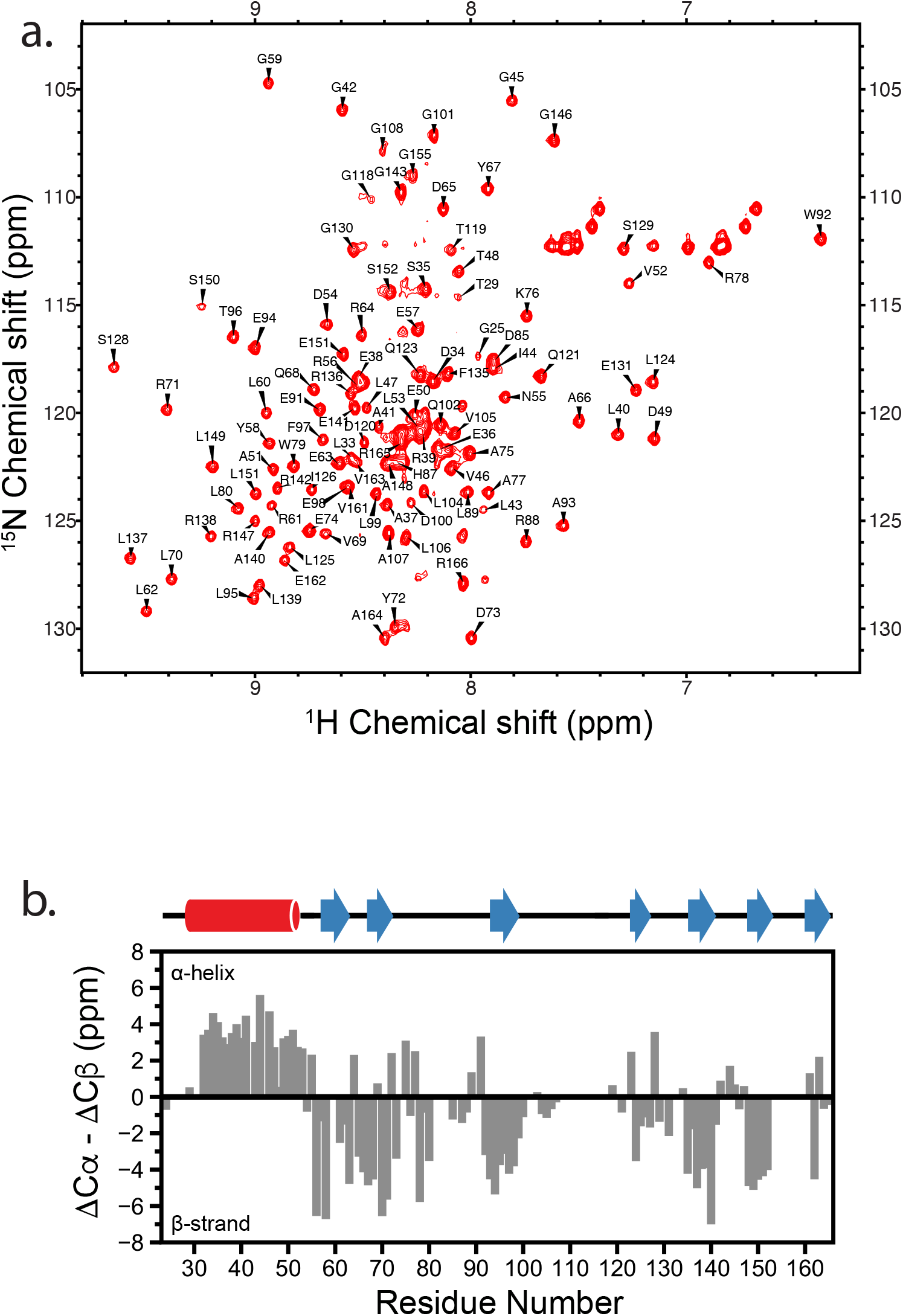
XcpHp NMR assignments and secondary structure determination. a. XcpHp assignments by solution NMR. 2D 1H-15N HSQC spectrum showing amino acid assignments for XcpHp. NMR experiments were performed at 37°C using a sample of 500 µM 13C-15N labeled XcpHp in NMR buffers. b. Secondary structure analysis of XcpHp. Secondary chemical shifts were calculated for XcpHp using alpha and beta carbon chemical shifts obtained after backbone sequential assignments. Stretches of amino acids with positive values indicate the presence of helix while negative values correspond to strands. Amino acids not assigned or overlapped are represented by a value of zero. Secondary structure elements in the XcpHp model structures are shown above.

We employed these assignments to experimentally determine the secondary structure elements within XcpH_p_ (**Figure 2b**). Furthermore, we generated two 3D homology models of XcpH_p_ using PDB 2KNQ and 2QV8^39^. Each 3D model matched our experimental secondary structure assignments well (not shown).

A subset of XcpH_p_ amino acids, in particular lysines, could not be assigned in our NMR spectra. The inability to assign these lysines implies a flexible region of the protein backbone. We tested the role of these residues in secretion by creating an in-frame deletion of amino acids 108-117, which form a long excursion between *β*3 and *β*4, resembling an old-fashioned mouse trap spring in one homology model (**Supplemental Figure 1**). We found that trans complementation with the *xcpH*_*Δ34*_ deletion restores LasB secretion and activity to the *ΔxcpH* strain (**Figure 1b, c**). Thus residues 108-117 are not required for LasB secretion, and therefore an disorder to order transition of this region cannot be required for secretion of LasB either.

#### Specific sites of spectral changes along the XcpH_p_ helix suggest binding interface: Chemical Shift Perturbation

With XcpH_p_ chemical shifts in hand, we performed chemical shift perturbation (CSP) analysis to identify structural regions impacted in XcpH_p_ upon binding of its partner XcpJ_p_. It is expected that resonances from residues close to the protein-protein interface will be shifted by the presence of the binding partner. Thus, a sample of 100 µM ^15^N uniformly labeled XcpH_p_ was titrated with increasing amounts of XcpJ_p_ and the 2D ^1^H-^15^N HSQC spectrum was collected. XcpJ_p_ concentrations above 80 µM could not be used since at that concentration there was a significant signal intensity loss due to the large size of the complex. Significant CSPs were observed in multiple places in the amino acid sequence (**Figure 3a)**. By mapping these CSPs on a structural model of XcpH_p_, we observed one subset of amino acid residues located close to the tip of the alpha helical spine of XcpH_p_ (residues G42 to G59 in the helix, and G130 and G131 on the nearby β4-β5 loop) that are perturbed by the presence of XcpJ_p_ (**Figure 3b)**. In addition, there is a second set of residues with significant CSP located on the C-terminal *β* sheet of XcpH_p_ (residues F135 to R138 in *β*5, L149 and S153 in *β*6 and E162 and A164 in *β*7). These additional perturbed sites may indicate a secondary point of interaction with XcpJ_p_, or changes in XcpH_p_ propagated through its structure after binding to XcpJ_p_. Global smaller amplitude changes across much of XcpH_p_ suggest that XcpH_p_:J_p_ interaction subtly affects dynamics or the structure of the entire XcpH_p_ molecule.

**Fig 3.**
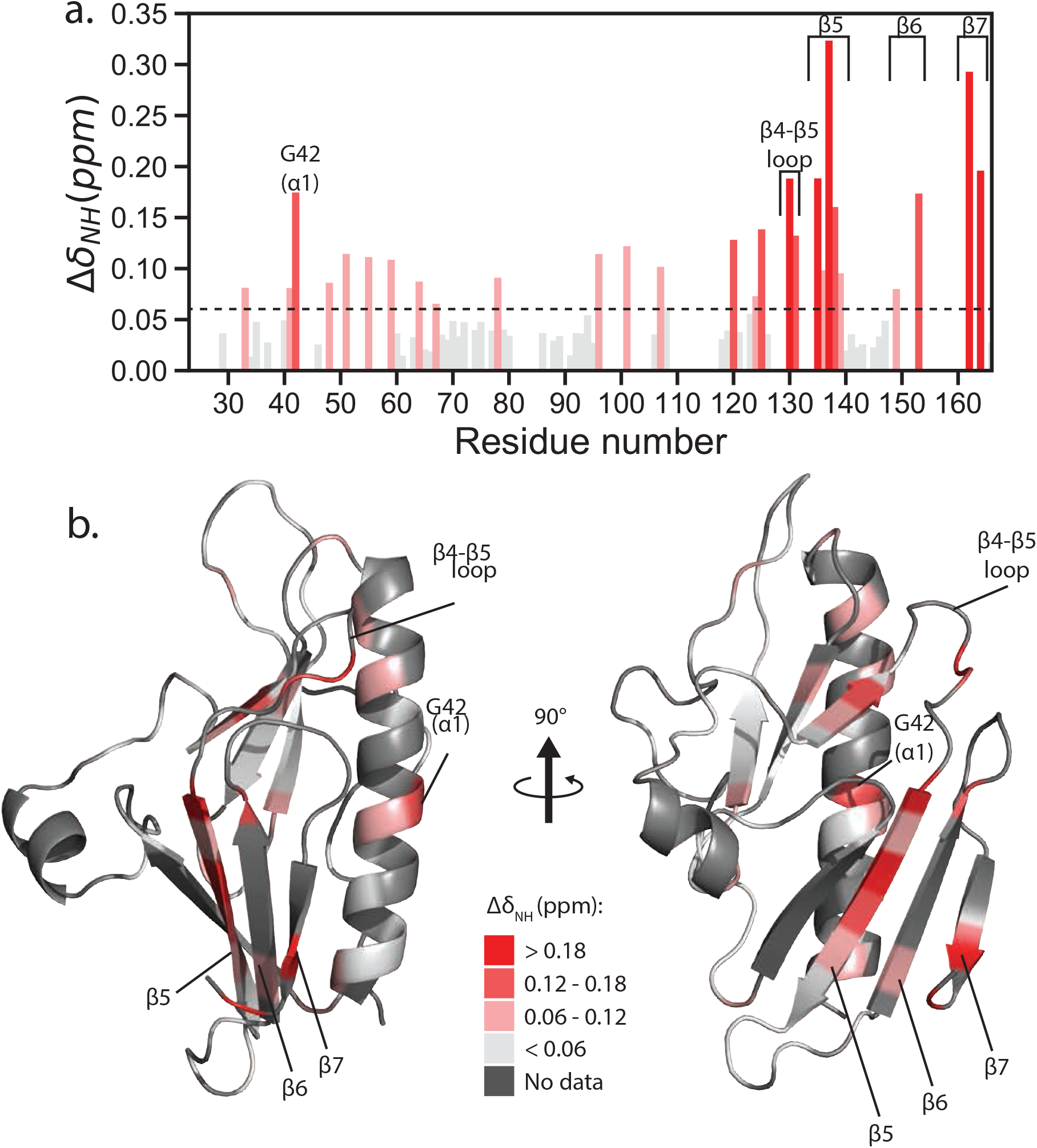
XcpHp chemical shift perturbations due to interaction with XcpJp. a. Plot of CSPs calculated for XcpHp amino acids. Each value was calculated using NH chemical shift differences obtained from the XcpHp spectrum collected without XcpJp and or with 80 µM XcpJp. CSP values above 0.06 were considered significant. Amino acids presented with value zero correspond to residues unassigned or overlapped. b. CSP values were mapped onto an XcpHp model based on E. coli GspH (PDB: 2KNQ64). Non-significant changes are shown in white, while more significant values are depicted in a gradient of red. Amino acids with no data are shown in grey.

### Site specific cysteine incorporation in XcpJ_p_ provides XcpH_p_:J_p_interaction sites

#### Cysteine cross-linking

The knowledge that a significant structural interaction interface for the major pseudopilins is the packing of their *α*1 helices^15^, combined with our CSP results implicating the XcpH *α*1C in the XcpH_p_:J_p_ interaction, led us to hypothesize that *α*1C:*α*1C packing forms a major interaction between these two minor pseudopilins. We introduced cysteines at surface-exposed *α*1C positions in XcpH_p_ (V46, D49, L53) and XcpJ_p_ (R46, R53) and mixed purified proteins under oxidizing conditions. In every instance, homodimers are formed (**Figure 4**). Heterodimers between XcpH_p_ and XcpJ_p_ were also observed in some prominent cases, most notably for XcpJ_R46C_:H_L53C_ (**Figure 4**). Heterodimers were weak or absent when XcpH_pV46C_ was one of the partners. Although these results provide no quantitative measure of affinity or specificity, they served as motivation to use XcpJ_p_ *α*1C cysteine variants as probes of specific sites of interaction in our NMR platform.

**Fig 4.**
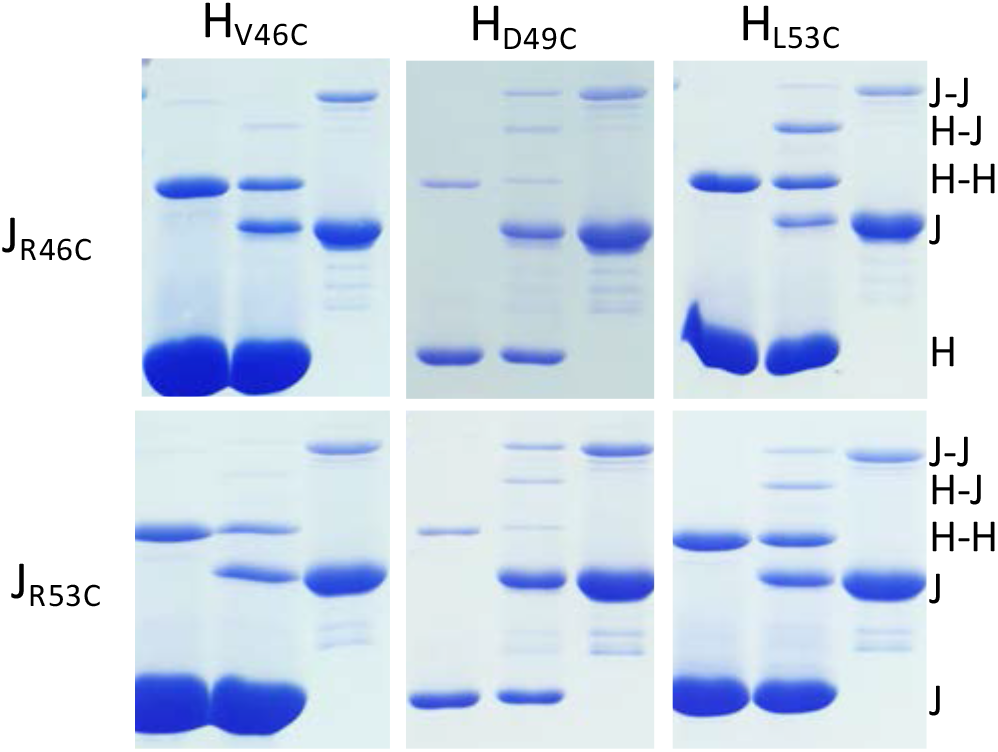
Cysteine cross linking suggests XcpHp:XcpJp 1C packing. Cysteine substitutions were made in indicated positions within XcpHp (by column) and in XcpJp (by row). Purified proteins were incubated under oxidizing conditions and visualized by Coomassie blue staining of non-reducing SDS PAGE. In each of the six pairwise combinations, Lane 1 is XcpHp alone, Lane 2 is XcpHp+XcpJp, and Lane 3 is XcpJp alone. Monomer, dimer, and heterodimer sizes are indicated along the right-hand side for all three gels in each row.

#### Paramagnetic relaxation enhancement

To identify local interactions and specific amino acid involvements in the XcpH:XcpJ interface, we performed paramagnetic relaxation enhancement (PRE) experiments guided by cysteine cross-linking results. In this approach, the effect of a specifically incorporated spin-label (1-Oxyl-2,2,5,5-tetramethylpyrroline-3-methyl) methanethiosulfonate (MTSL) within XcpJ_p_ on the ^15^N XcpH_p_ spectrum is observed; in particular, XcpH_p_ residues in proximity to the spin label on XcpJ_p_ will experience a decrease in signal intensity. XcpJ_p_ cysteine variants R46C and R53C were spin-labeled, as these positions in XcpJ_p_ were implicated in cysteine cross-linking experiments described above to participate in an XcpH α1C - XcpJ α1C packing interaction **(Figure 4)**. In addition, XcpH_pT178C_ and XcpH_pE180C_ were created and used for spin-labeling; these amino acids are on the face of XcpJ opposite α1C, where we did not expect any change in signal from spin label incorporation. We chose to perform PRE experiments using a 10:3 molar ratio of ^15^N-U XcpH_p_ to spin-labeled XcpJ_p_ variants. Although a 1:1 ratio would in theory have provided more quantitative results, for this protein pair, it would have led to signal loss as seen in the previously described titration.

The spin label MTSL at positions 46 or 53 within XcpJ_p_ has a marked effect on XcpH_p_ signal intensities (**Figure 5a**). XcpJ_pR46C_-MTSL induced a signal reduction on XcpH_p_ α1C (residues 42 to 54), with a maximum effect on D49. In addition, residues located in the loop between XcpH_p_ β4 and β5 are also affected (G130 and E131) (**Figure 5**), in agreement with CSP results. XcpJ_pR53C_-MTSL had an effect on similar amino acid residues. Within XcpH_p_ α1C the overall signal intensity decrease was slightly less compared to the C46 label, but interestingly the effect was shifted up the helix with the residue most greatly affected by the paramagnetic label, XcpH_p_-N55, located at the tip of XcpH_p_ (**Figure 5**). No significant effect was observed on XcpH_p_ when XcpJ_p_ was labeled at positions T178C or E180C (**Supplemental Figure 2**). Thus, the PRE experiment clearly supports that the interface between XcpH_p_ and XcpJ_p_ is mediated by α1C interactions with involvement of residues in the β4-β5 loop of XcpH. In addition, the difference in signal intensity decrease and the shift on the most affected residue towards the tip of XcpH_p_ when the MTSL label is at XcpJ position 53 as opposed to 46 suggest that the helical register is organized with XcpJ_p_ above XcpH_p_ in the heterodimer.

**Fig 5.**
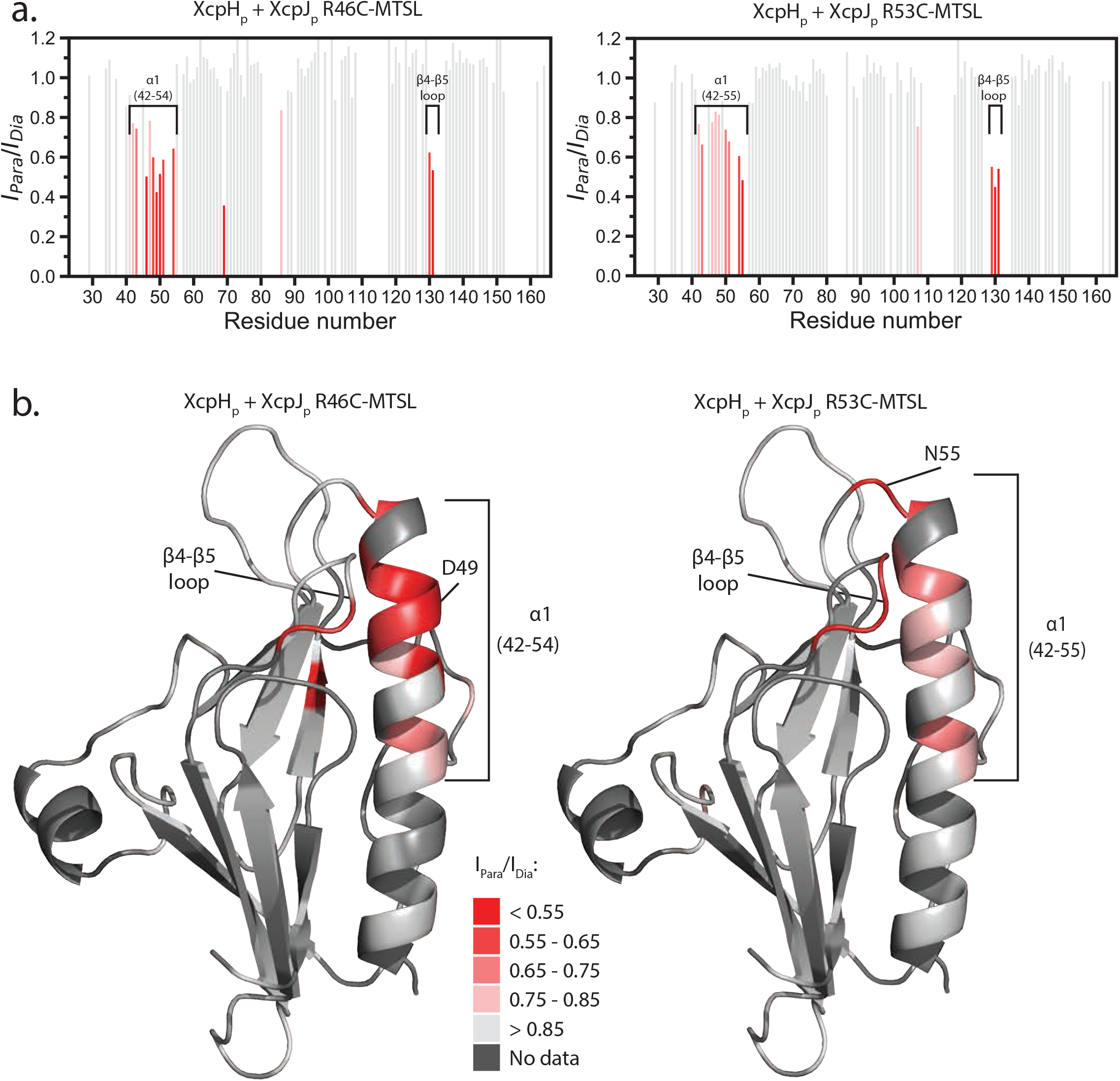
PRE experiments of XcpHp in the presence of MTSL labeled XcpJp. a. PRE effect of the two alpha1C MTSL-labeled XcpJp cysteine mutants on the XcpHp spectra. Spectra were collected using 80 µM 15N XcpHp and 24 µM XcpJpR46C-MTSL or XcpJpR53C-MTSL. Significant signal intensity changes are designated with a gradient of red. All amino acids with displayed data have an intensity ratio above zero, while residues with no assignments or overlapped have values of zero. b. PRE effects observed with XcpJpR46C-MTSL or XcpJpR53C-MTSL were mapped onto the structural model of XcpHp.

### Cross-linking Mass Spectrometry using acidic cross links *via* the DMTMM activating reagent

In order to extend our site-specific information from engineered disulfide bonds and PRE measurements, we applied a recently developed methodology for crosslinking of acidic side chains within a defined distance, followed by mass spectrometry identification of cross-linked peptides^50^. We established our pipeline using a 1:1 molar ratio of XcpH_p_ and XcpJ_p_ (Supplemental Methods and **Supplemental Figure 3**). Previous work has established that XcpJ_p_ is indispensable for XcpH_p_ involvement in the quaternary complex, which was the motivation for using this pair in our NMR experiments also^27,28^. Subsequently, we interrogated the complete quaternary complex of XcpH_p_I_p_J_p_K_p_, known already to form *in vitro* a quaternary complex with a molar ratio of 1:1:1:1^28^. These latter experiments confirmed the set of acidic crosslinks between XcpH_p_ and XcpJ_p_ are the same in the heterodimer as in the heterotetramer, and thus we focus here on the quaternary complex results.

The four minor pseudopilin soluble constructs XcpH_p,_ XcpI_p,_ XcpJ_p_ and XcpK_p_ contain 25, 14, 36 and 42 acidic residues, respectively, which are distributed along their amino acid primary sequences. Thus the use of adipic acid dihydrazide (ADH) coupled to 4-(4,6-dimethoxy-1,3,5-triazin-2-yl)-4-methyl-morpholinium chloride (DMTMM) as a coupling reagent appeared promising. DMTMM activates carboxylic acid functionalities, usually poorly reactive, allowing coupling of the crosslinking reagent ADH to the extended amino acid side chains with high efficiency^50^. When added in large excess to a freshly prepared aqueous solution containing a 1:1:1:1 molar ratio of XcpH_p,_ XcpI_p,_ XcpJ_p_ and XcpK_p_, DMTMM/ADH reagents formed covalent bonds (**Figure 6a**), which connect proximal acidic functionalities with a maximum linking distance of 20-25 Å^51^. The most likely tetramer band observed on SDS-PAGE of this mixture was excised and analyzed by a classical proteomic bottom-up approach to validate the presence of all four proteins. Each protein was identified unambiguously, with a sequence coverage of 92% for XcpH_p_ (61 peptides), 78% for XcpI_p_ (31 peptides), 86% for XcpJ_p_ (71 peptides) and 91% for XcpK_p_ (146 peptides).

**Fig 6.**
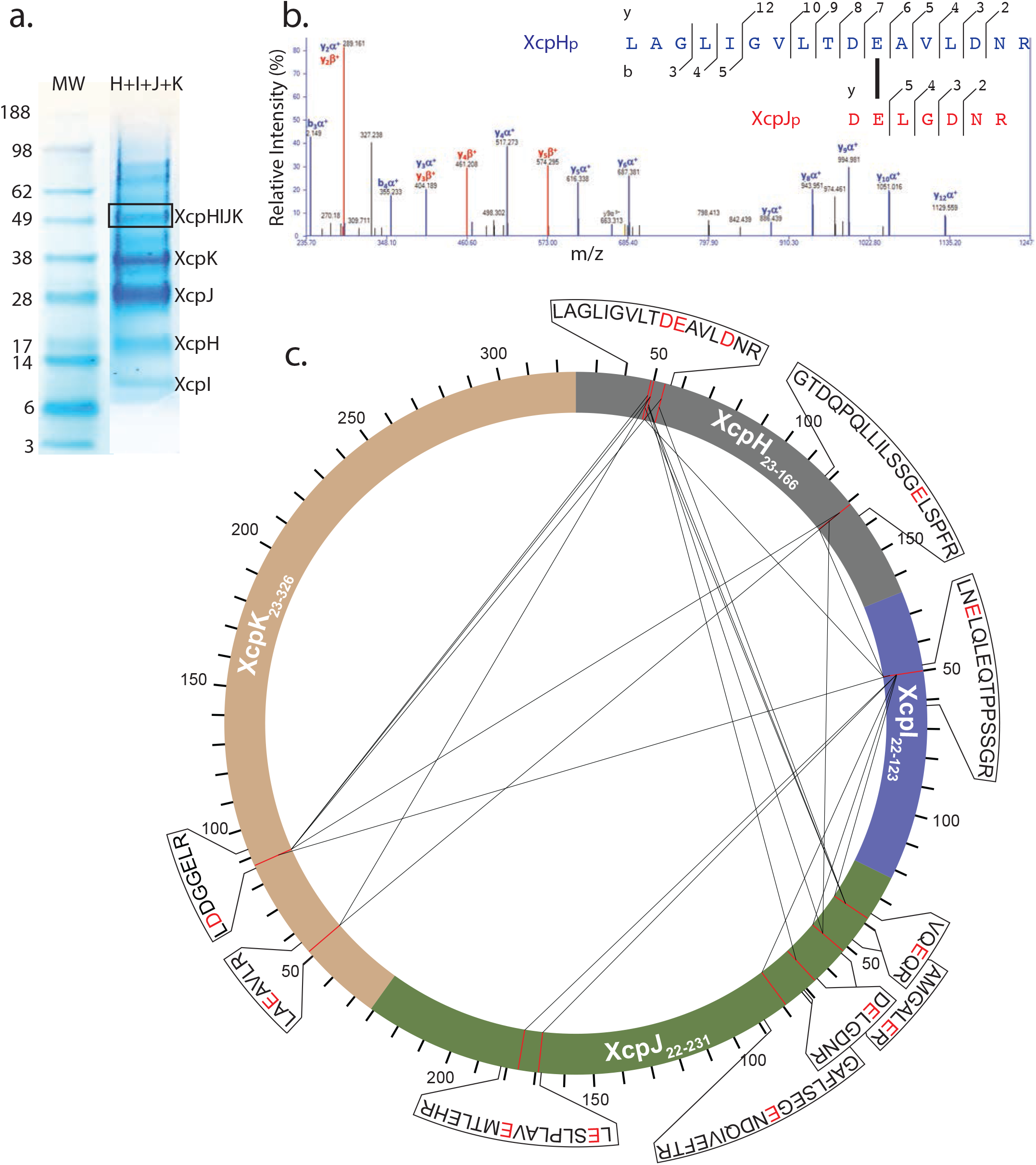
Acidic crosslinking of XcpHpIpJpKp soluble complex. a. SDS-PAGE of the cross-linked proteins with a box around the band identified by the mean of proteomics as the cross-linked XcpHpIpJpKp tetramer. b. MS/MS spectrum of a particularly relevant cross-link pair between XcpHp and XcpJp. The spectrum shows intense ions identified as y- and b-type ions, allowing an unambiguous identification of the two partners (mass accuracy for the parent < 2ppm and for the fragment <10ppm). c. Circular representation of all significant cross-linked pairs detected in the tetramer.

Due to the long length of the linkers, these cross links provide complex and valuable information to constrain a model of the quaternary structure of the assembled XcpH_p_I_p_J_p_K_p_ complex. Most revealing for our goals of understanding the placement of XcpH within the pseudopilus quaternary complex were crosslinks between glutamate or aspartate side chains of XcpH_p_ with any of the other three soluble pseudopilin subunits in the complex. Many inter-subunit crosslinks between XcpH_p_ and XcpJ_p_ were identified between α1C of XcpH_p_, in particular within the peptide LAGLIGVLTDEAVLDNR, and the several peptides that together span the analogous α1C of XcpJ_p_ and the *β* hairpin that follows it (**Figures 6b, 6c**). An example MS/MS spectrum reveals intense peaks defining the connection between the two peptides (**Figure 6b**), with both chains unambiguously characterized by many fragment ions distributed along the peptide sequences. Moreover, the large mass difference observed between y_6_*α*^+^ and y_7_*α*^+^ characterizes precisely the position of crosslinking. The same α1C region of XcpH_p_ also cross links with two regions of XcpK_p_ and one region of XcpI_p_ therefore suggesting they are in physical proximity (**Figure 6c, Table S3**). Interestingly, an acidic residue located in the XcpH_p_ β_4_β_5_ loop shown by PRE analysis to be in close proximity with the α1C region of XcpJ_p_ (**Figure 5b**) is also involved in cross links with XcpJ_p_, XcpI_p_ and XcpK_p_ (**Figure 6c, Table S3**).

Crosslinks between XcpI_p_ and XcpJ_p_ as well as a single crosslink identified between XcpI_p_ and XcpK_p_ were all in agreement with the crystallographically determined structures of GspI_p_J_p_K_p_ and XcpI_p_J_p_K_p_. Moreover, intrasubunit crosslinks agreed with known protein 3D structures (data not shown). This rich library of crosslinks between amino acid side chains within the quaternary complex provides strong constraints for modeling of a three-dimensional structure of the XcpH_p_I_p_J_p_K_p_ complex.

### Integrating experimental data to build quaternary complex model

#### XcpH_p_I_p_J_p_K_p_ complex

Data from spin labelling studies and acidic cross linking mass spectrometry were used as restraints on the interaction of XcpH_p_ with XcpI_p_J_p_K_p_, to model the quaternary complex using HADDOCK software for protein-protein docking. Docking used the structure of the XcpI_p_J_p_K_p_ ternary complex (PDB: 5VTM), the XcpH_p_ homology model, PRE experimental results and acidic cross-linking interactions as input (**Supplemental Tables 3 & 4**). The best model obtained is in good agreement with these experimental restraints (**Figure 7**). In this model, the α1C of XcpH_p_ is in direct contact with that of XcpJ_p_, and runs approximately parallel to the helical bundle in the XcpI_p_J_p_K_p_ complex (**Figure 7**). XcpH_p_ interaction with the ternary complex is maintained by a large network of electrostatic interactions and hydrogen bonds (**Figure 7**). One group of contacts corresponds to an electrostatic interaction between helices, in particular including a salt bridge between side chain XcpH_p_ D49 and XcpJ_p_ R42 (possibly also to XcpJ_p_ R46, given the rotamers available to Arg side chains and the limitations of resolution of this model) as well as numerous main chain and side chain H-bonds. A second group comprises β-strand to β-strand contacts. In this interaction, XcpH_p_ loop β6-β7 G155 and F156 backbone carboxyl groups form hydrogen bonds with XcpJ_p_ β7 R131 side chain and Q133 main chain, respectively. Other contacts observed between XcpH_p_ and XcpJ_p_ correspond to: 1) XcpH_p_ loop α1-β1 and XcpJ_p_ loop β4-β5 (N55-R92), 2) XcpH_p_ loop β4-β5 and XcpJ_p_ loop β4-β5 (S129-R92) (**Figure 7**). Remarkably, although no data from the chemical shift perturbation experiments were explicitly included in the HADDOCK restraints, many of the interactions in the model are recapitulated as chemical shift perturbations, for example the very high signal interaction within XcpH_p_ β7 is explained by its close approach to XcpJ_p_.

**Fig 7.**
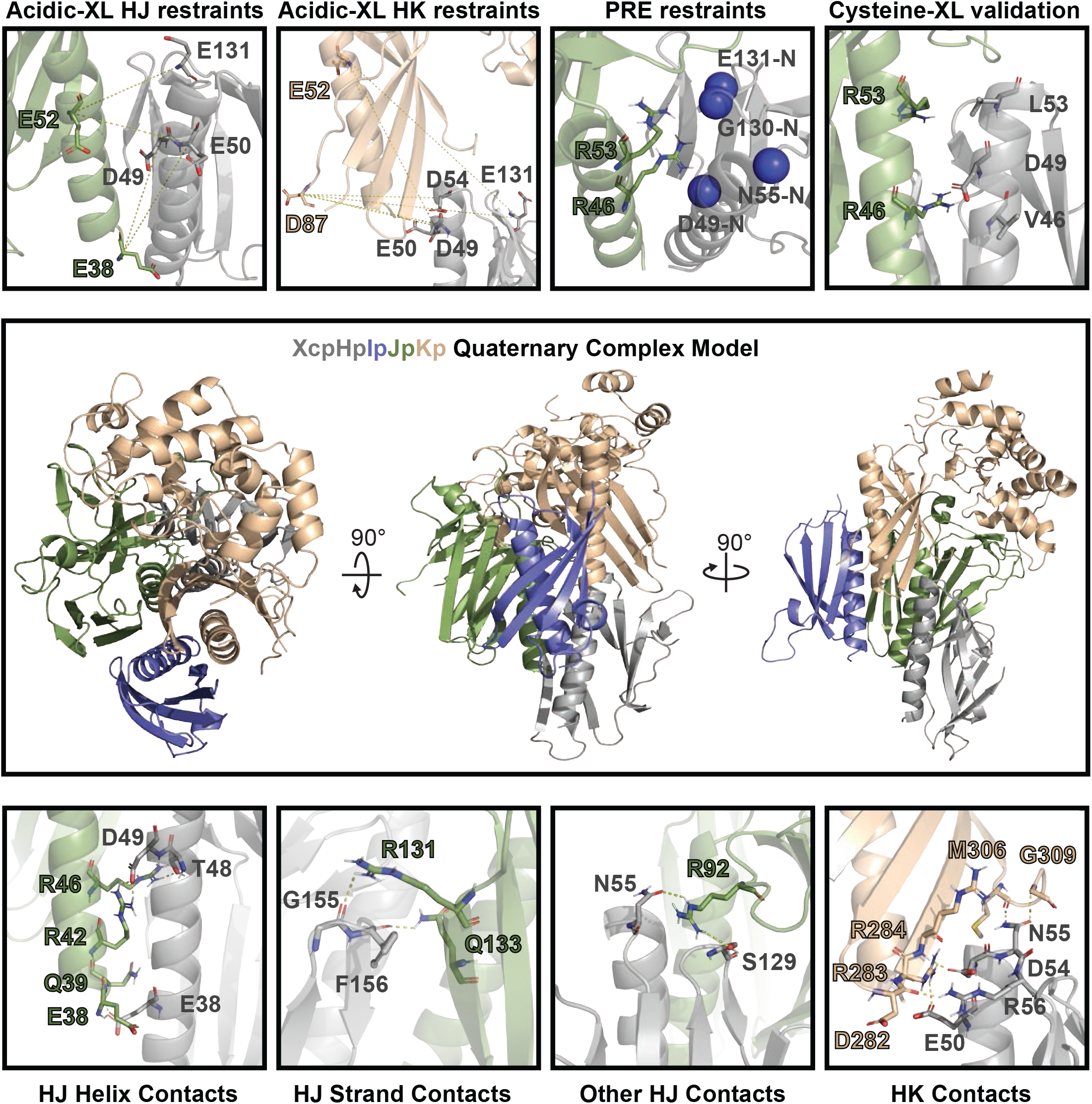
3D model of XcpHpIpJpKp soluble complex. A model of the XcpHpIpJpKp quaternary complex (XcpHp gray, XcpIp blue, XcpJp green and XcpKp wheat) was obtained via HADDOCK docking. Acidic cross-linking derived restraints are shown in yellow dotted lines. In the case of PRE restraints, XcpJp spin labeled residues R46 and R53 are shown as sticks while most affected XcpHp residues N55 and D49 are represented as blue spheres at the amide nitrogen. XcpHpJp residues forming salt bridges or hydrogen bonds are shown in stick model connected by yellow dotted lines. Residues from XcpHp and XcpJp are labeled in dark grey and dark green respectively

The use of acidic cross-linking restraints for XcpH_p_ with the two other subunits from the quaternary complex disambiguates the XcpH_p_ position *vis a vis* these proteins, for example the crosslinks to XcpK_p_ help to position the tip of XcpH_p_ α1C close to the XcpK_p_ β-sheet, allowing additional contacts that stabilize the complex. These include XcpH_p_-XcpK_p_ residues E50-R283, D54-R283, R56-D282, R56-R284 and N55-G309 (**Figure 7**). We were able to further support this quaternary structure by analyzing the co-evolution of residues in XcpH_p_ and XcpJ_p_. (**Supplemental Figure 4**). The analysis similarly points to XcpH polar amino acids at the tip of *α*1C (D49, E57) and in the *β*4-*β*5 loop (S128, S129, E131) and pairs them with two sets of XcpJ co-evolved amino acids located within a large flap at the tip of XcpJ (R92 and R99, forming electrostatic interactions), or in the vicinity of *α*1C (V45, R189 and W107, potentially contributing to packing geometry) (**Supplemental Figure 4**). This co-evolution result adds credence to the robustness and physiological relevance of our XcpH:XcpJ interaction model. Altogether, we present a model of the XcpH_p_I_p_J_p_K_p_ quaternary complex supported by experimental evidence, which shows the interaction of XcpH with XcpJ and XcpK is maintained by several electrostatic interactions.

#### Filament model

The quaternary complex presented here reveals the interaction of the minor pseudopilin soluble domains, and in particular places XcpH_p_ in this ensemble. However, these proteins also interact via their missing hydrophobic *α* helices (α1N) in the biological context of the pseudopilus. Thus, we used available experimental data and chemically reasonable restraints to model the complete Type II secretion system pseudopilus. For that purpose, hydrophobic α1N were added to the XcpH_p_I_p_J_p_K_p_ proteins in the soluble quaternary complex model. In addition, the major pseudopilin XcpG filament was modeled using as template the structure of PulG filament (PDB: 5WDA)^15^. Then, the quaternary complex was added to the tip of the XcpG filament by aligning XcpH to the first XcpG unit. Finally, the whole model was minimized using PyRosetta^52^. The best model was selected based on the lowest RMSD with respect to the HADDOCK model (**Figure 8**). An important feature included as a restraint in calculating this model was the salt bridge between N-terminal amino groups and the carboxylic acid of E5 of each preceding unit in the filament, including XcpHIJK at the tip of the pseudopilus. This contact neutralizes these charges in the transmembrane helices as seen in other filament structures such as *P. aeruginosa* PAK pilus (PDB 5VXY). As expected, E5-F1 contacts continue into the minor pseudopilin complex in the following sequence: XcpG-XcpH-XcpJ-XcpI-XcpK. This arrangement maintains the right-handed nature of the major pseudopilin filament structure. In the filament model, the average distance between E5 carboxyl group and F1 amino group is 3.58 ± 0.34 Å between adjacent subunits. In addition, Cisneros *et al*.^37^ showed the close contact between the residues 16 and 10 of neighboring subunits (PulJ-PulI and PulI-PulK). Even though these contacts were not used as restraints during modelling, the current model reproduces them between the minor and major pseudopilins throughout the filament (**Figure 8** and data not shown). Thus, the C*α*-C*α* distance from residue 10 to 16 of the neighbouring subunits in the PulG filament is 8.5 Å, in the model that distance is on average 8.9 ± 1.4 Å. Strikingly, to maintain the restraint contacts, in particular the E5 to F1 salt bridge between XcpH and XcpJ, it was necessary to allow XcpH α1N between residues 20 and 26 to unravel. Although there is no other experimental evidence that the XcpH helix adopts this melted secondary structure, the equivalent amino acids in the major T2SS pseudopilin and the major T4P pilin are strikingly extended in high resolution filament models^15,53^. Additionally, the presence of a glycine at XcpH position 25 supports this conclusion, since glycine and proline residues destabilize *α* helices.

**Fig 8.**
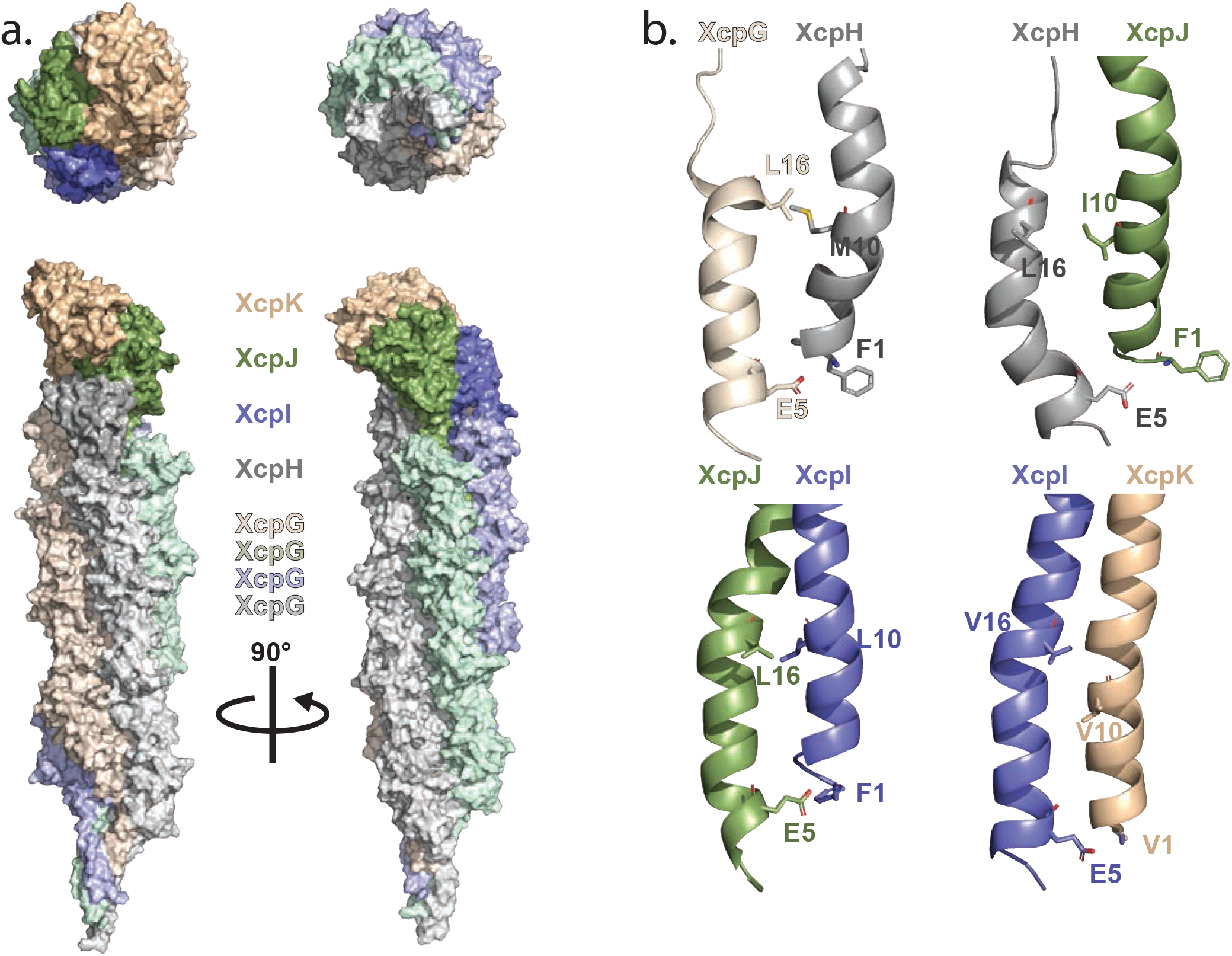
Model of Type II secretion system pseudopilus. a. Model of XcpHIJK tip complex and XcpG filament shown as a solvent-accessible surface. (above) Top and bottom views of the models (below) Side views of the pseudopilus model. Color coding of each XcpG protofilament is a lighter shade of the respective tip subunit color from whence it originates. b. Transmembrane alpha1N contacts between neighboring subunits include canonical F1-E5 salt bridge (included as restraint during minimization), and hydrophobic residues at position 16 and 10 previously described (37) but importantly not used as a modeling restraint. Images were generated using Pymol software.

Another feature observed in the filament model, specifically related to the minor pseudopilin tip, is the capping of each XcpG protofilament by a minor pseudopilin and the off-center placement of XcpK. If the major pseudopilin filament is separated into 4 protofilaments, each one is capped by a minor pseudopilin (**Figure 8**). The exception to this is XcpJ; although the XcpJ globular domain is directly above one protofilament, a small gap exists between XcpJ and the XcpG unit directly below it. An additional consequence of capping one XcpG protofilament with XcpH is that the XcpK globular domain protrudes beyond the major pseudopilin filament diameter, giving the appearance of a hook-like structure to the filament tip (**Figure 8)**.

While our model cannot be considered an atomic structure of the Type II secretion pseudopilus, it provides a valuable path to testable predictions such as the importance of XcpH helix unravelling, possible substrate binding sites available on the filament tip, the presence of dynamic contacts between XcpG and minor pseudopilins, and the importance of the XcpK globular domain for secretion.

## DISCUSSION

Study of dynamic interactions prioritized by binding affinity is challenging but we have used chemical, structural, microbiological, and computational approaches to describe the complete ultrastructure of the T2SS pseudopilus and in particular to propose a specific and testable model for the location of the adapter unit XcpH (GspH) between XcpIJK and the filament formed by XcpG. Addition of XcpH would be the next step in a dynamic chronological pathway following XcpIJK association at the periplasmic membrane. Subsequently, XcpG addition absolutely requires the *α*1N helices, as there is no interaction between the soluble domains of XcpH and XcpG (**Supplemental Figure 5**), nor indeed between the soluble domain of XcpG and the soluble domain of any of the quaternary complex soluble domains individually^27^. We have amassed independent and complementary data from several techniques in this integrative study. In particular, the pioneering methodologies in acidic crosslinking represent an important technical breakthrough in the experimental approach to studying protein:protein interactions in multipartner complexes.

This study provides a rationale for the observation that at least in the *P. aeruginosa* Xcp T2SS, XcpH is required for secretion. It serves as the adapter that connects the regular helical assembly of XcpG subunits to the asymmetric complex that catalyzes initiation of the filament and interacts with substrates and other components of the secretion system. It may be possible that XcpH serves other as yet unverified roles in the system as well. For example, given its very strong structural homology to the major XcpG subunit, XcpH might occasionally integrate into the filament and thereby destabilize it, possibly initiating disassembly of the pseudopilus^31^.

Previous interaction studies using surface plasmon resonance^27^ testing pairwise interactions of the soluble domains of XcpG, H, I, J and K as well as our current NMR results (**Supplemental Figure 5**) reveal that XcpH interacts exclusively with XcpJ. In addition, the binding of XcpH to XcpJ is more efficient when XcpJ is engaged in the XcpIJK ternary complex (Figures 2 & 3 in ^27^). These data suggested that conformational changes in XcpJ upon binding to XcpI and XcpK create a more favorable docking platform for XcpH and/or favor the downward addition of XcpH to the ternary complex over that of XcpG when both are available within the inner membrane. Our current data supporting many pairwise interactions among the quaternary complex proteins, including between XcpH and each of the other three, fit well with the hypothesis that alone the soluble domains have weak and transient interactions with each other which in some cases are below detection but that within the full ensemble, these interactions are revealed. Our model furthermore accentuates that the central *α* helices are, not surprisingly, fundamentally important for more stable interactions. Future experiments will rely on the *α*1N helical tails being part of the interactions.

Based on the near atomic resolution of the XcpQ secretin^12^ and taking into account that the pseudopilus tip physically interacts with the secretin^33^, a satisfying outcome of our data-driven modeling of the complete pseudopilus is the observation that the filament (∼6 nm) docks readily into the periplasm-facing vestibule (∼8 nm)^12^ of the XcpD secretin (**Supplemental Figure 6**). Moreover, the positioning of the Xcp pseudopilus plus secretin complex into the cryo-tomographic map of the complete T2SS in its natural context, the bacterial envelope^23^, indicates that 8 to 16 pseudopilin subunits in addition to the tip complex are required to span the periplasm between the IM and the entry into, or the internal periplasmic gate of, the secretin interior, respectively (**Supplemental Figure 6**).

The results presented here provide insights into not only the *P. aeruginosa* T2SS but indeed into the dozens of T2SS recovered in proteobacteria with similar genome content of five pre-pseudopilins equivalent to XcpG, H, I, J, and K^2^. The conservation of amino acid sequences of these proteins across species implies their protein:protein interaction interfaces are also conserved, and coevolved as shown here for the XcpH_p_:XcpJ_p_ interface. By extension, both the minor pseudopilins of the T2SS and the minor pilins of the T4P can be expected to form a similar quaternary complex in which an adapter minor (pseudo)pilin subunit sets the stage for downward addition of the major (pseudo)pilin subunit to form the extended filament by interacting via its periplasmic domain with the three upper minor subunits and via its unraveled *α*1C with the first major subunit. Correspondingly, the set of *E. coli* T4P minor pilins restore assembly of the major T2SS pseudopilin PulG in absence of minor pseudopilins PulHJIK^37^. An open question is whether the secretion ATPase motor protein, GspE, is required for insertion and/or *α*1C unraveling in this adapter minor pilin at the junction between the tip complex and the filament itself.

## ONLINE MATERIALS AND METHODS

### Gene and protein nomenclature

Here we have extended the general T2SS nomenclature to the *P. aeruginosa* Xcp T2SS components. Thus, former XcpT, U, V, W and X proteins are now labelled XcpG, H, I, J and K in agreement with their homologs in other T2SSs. All pseudopilin soluble domains (deleted for their N-terminal hydrophobic domains (α1N)) used in this study are designated with a subscript p indicating periplasmic. According to the standard in the field, polypeptides are numbered with amino acid 1 as the first residue in the mature pseudopilin following prepilin peptidase cleavage.

### Bacterial strains and plasmids

*Escherichia coli* K-12 DH5α (laboratory collection) and BL21(DE3) pLysS (laboratory collection) were used for cloning procedures and soluble protein production, respectively. *Pseudomonas aeruginosa* PAO1 wild type (laboratory collection), PAO1 *ΔxcpH* (this study), *ΔxcpI* (this study), *ΔxcpJ*^43^, and *ΔxcpK*^26^ or PAO1 *Δxcp* (also called DZQ40)^54^ strains were used for *in vivo* complementation assay. Construction of the *xcpH* and *xcpI P. aeruginosa* PAO1 deletion strains was performed as described previously^16^ using the pKN-*Δ*H and pKN-*Δ*I mutator plasmids (**Table S1)**.

Plasmids and oligonucleotides used in this study are listed in **Table S1** and **Table S2**, respectively. Site directed mutagenesis was performed using quick change technology (Stratagen) or inverse PCR. The gene encoding the XcpH_p_ protein deleted for its mouse trap domain was cloned into the pJN105 vector using SLIC technology^55^. Plasmid pET-XcpH_p_ was constructed following the strategy used by Durand et al.^16^ to construct pET-XcpG_p_. All constructs have been verified by DNA sequencing.

### LasB secretion and protease activity on plates

Preparation of culture supernatants from *P. aeruginosa* for analysis of secreted proteins has been described^56^. Gel analysis was standardized so that the volume of supernatant equivalent to 2 OD_600_ units of culture was used for each sample (**Figures 1a, 1c**). Protease activity was tested by spotting 5 µl of bacterial culture grown in LB to early stationary phase on TSA plates containing 1.5% lyophilized milk and appropriate antibiotics, followed by 14 h incubation at 30°C.

### Protein production and purification

Production and purification of XcpH_p_ and XcpJ_p_ with or without cysteine substitutions for NMR purposes (except XcpG_p_/H_p_ NMR CSP, see Supplemental Material) was carried out with minor modifications from published procedures^32^. Expression was performed in BL21(DE3) pLysS *E. coli* carrying the plasmids pETG-20A-XcpH_p_ or pETG-20A-XcpJ_p_. For ^13^C and ^15^N uniform protein labeling, cells were grown in 3 L of LB at 37°C to OD_600_ ∼0.6. Cells were collected and washed with M9 media without glucose or ammonium chloride to remove remaining LB media. After washing, cells were transferred to 1 L M9 media supplemented with 1 g of ^15^N-U ammonium chloride and 2 g of ^13^C-U glucose (Cambridge Isotope Laboratories) and incubated at 25°C for 2 hours before inducing with 1 mM IPTG overnight. Purification of XcpH_p_ proceeded using a combination of nickel affinity and size exclusion chromatography. The cell pellet was resuspended in Buffer A (50 mM tris-HCl buffer pH 8.0 containing 300 mM NaCl) and lysed using a French press. Supernatant from lysate clarified by centrifugation at 25,000 x G for 30 min. was loaded into a 5 mL Ni-NTA column (Quiagen) and washed with 100 mL of Buffer A containing 50 mM imidazole. Protein was eluted from the column using Buffer A with 500 mM imidazole. Peak fractions containing target protein were pooled and treated with TEV protease (purified in house) added to a final concentration of ∼40 µg/mL and the sample was dialyzed overnight at 4°C against Buffer A to remove imidazole. Removal of TEV protease proceeded by loading samples into a 1 mL Ni-NTA column which was washed with the same buffer. Fractions containing XcpH_p_ were collected and concentrated for size exclusion chromatography in a Sephacryl S-100 HiPrep 16/60 column (GE). During this step, buffer was exchanged to NMR buffer (25 mM sodium phosphate buffer pH 6.5 containing 25 mM NaCl). XcpJ_p_ was expressed and purified using the identical protocol but without ^13^C and ^15^N labeling.

Production and purification of XcpH_p_, XcpI_p_, XcpJ_p_, XcpK_p_, XcpH_pV46C_, XcpH_pD49C_, XcpH_pL53C_, XcpJ_pR46C_ and XcpJ_pR53C_ for cysteine cross-linking (and cross-linking MS experiments was performed in BL21(DE3) pLysS *E. coli* carrying the corresponding pETG-20A plasmid (**Table S1)** as published.^27^

### In vitro cysteine cross-linking

40 µM of a single cysteine variant or two variant proteins in 1:1 molar ratio were incubated in Tris-HCl 50 mM, NaCl 100mM, pH 8 in a total volume of 500 µL. The mixture was supplemented with DTT (20 mM) to allow reduction of intra-chain disulfide bands formed during purification steps. The mixture was dialyzed against 300 mL of Tris-HCl 50 mM, NaCl 100 mM, pH 8 buffer for 2 h at RT to allow DTT removal and cysteine oxidation. To analyze disulfide bond formation, 75 µL of each reaction was mixed with 25 µL of Leammli loading buffer with or without reductant as appropriate. The samples were heated for 5 min at 95°C and then visualized by 12% Coomassie blue stained SDS-PAGE.

### Acidic Crosslinking of XcpH_p_ + XcpJ_p_ and XcpH_p_ + XcpI_p_ + XcpJ_p_ + XcpK_p_

#### Preparation of multimers

For assembling the dimer, 17µL of XcpH_p_ at 4 mg/mL were mixed to 25µL of XcpJ_p_ at 4 mg/ml, leading to a mixture of 4 nmol of each protein (molar ratio 1:1). For the tetramer, 17µL of XcpH_p_ (4 mg/mL), 12µL of XcpI_p_ (4 mg/mL), 25µL of XcpJ_p_ (4 mg/mL) and 34µL of XcpK_p_ (4 mg/mL) were mixed to yield a 4 nmol solution of each protein (molar ratio 1:1:1:1). Samples were dried under vacuum, and dissolved in 100µL of PBS buffer pH 7.4 to achieve a final concentration of each protein of 40µM. Proteins were kept at 25°C for 1 h to let the association of the dimer or tetramer occur.

#### Cross-linking reaction

To crosslink the dimer, 10µL of ADH (adipic acid dihydrazide) and 16µL of DMTMM ((4-(4,6-dimethoxy-1,3,5-triazin-2-yl)-4-methyl-morpholinium chloride) were added to the solution of XcpH_p_:J_p_ complex, reaching final concentrations of ADH and DMTMM equal to 46 mM (8 and 12.7 mg/mL, respectively). For the tetramer, 23 µL of ADH and 36 µL of DMTMM were added to the solution of XcpH_p_:I_p_:J_p_:K_p_ complex, reaching final concentrations of ADH and DMTMM equal to 81 mM (14.5 and 22.6 mg/mL, respectively). These concentrations were chosen to impose a molar excess of crosslinking reagent of more than 1,000. The mixture was incubated 2 h at 37°C under agitation (750 rpm) to allow intensive crosslinking reaction. The reaction was quenched by reagent removal using a ZebaSpinDesalting Column (0.5mL, 7k, Pierce), employed according to the manufacturer recommendations. The resulting sample was dried using a Speedvac system.

#### SDS-PAGE

Cross-linked species were imaged by classical SDS-PAGE analysis. The protein mixture was suspended in 170µL of 8M urea, leading to a global protein concentration of 48µM. 20µg of proteins (=10µL) were dried before being solubilized in 20µL of Laemmli buffer (Tris HCl pH 6.8 (65mM), SDS 2%, glycerol 20% and DTT 350mM) and a spatula tip of bromophenol blue. The sample was heated at 100°C for 5 min, and 10µL (∼10µg) were loaded on the gel (NuPageTM 4-12% Bis-Tris-Gel, 1.0 mm x 10 wells). The migration starts with a voltage of 200V applied for 40min (stacking), and then 150V for the separation. The electrophoresis system is switched-off when the migration blue line is localized at the extremity of the gel. The gel is fixed 3h in a bath composed of 50% ethanol, 47% water and 3% phosphoric acid, washed three times with ultrapure water, and finally incubated in water containing 34% methanol, 17% ammonium sulfate and 3% phosphoric acid. The coloration of the bands was made during 3 days with Coomassie blue G250 added at 360 mg/l in the solution. The gel was finally washed several times in pure water to remove the excess of Coomassie blue.

#### In-gel digestion

The bands corresponding to the XcpH_p_:XcpJ_p_ dimer or XcpH_p_:XcpI_p_:XcpJ_p_:XcpK_p_ tetramer, respectively, were excised, and cut into small pieces. Resulting pieces of gel were then washed by 50µL of 50mM NH_4_HCO_3_ then centrifuged (5min, 600 rpm) to remove the supernatant. The same step was repeated with 50µL of 50/50 (v/v) 50mM NH_4_HCO_3_/acetonitrile. The two previous steps were repeated twice. To ensure a good penetration of the enzymes into the pieces of gel, the latter were dehydrated twice with 50µL of pure acetonitrile removed by centrifugation (5 min, 600 rpm). The gel was then rehydrated at 0°C with 3µL of a solution containing 1/100 of Lys-C and 1/50 of trypsin in 50mM NH_4_HCO_3_. The digestion was performed at 37°C for 4 hr. Enzymatic activity was quenched by acidifying the medium using 25µL of 1% trifluroroacetic acid. Resulting peptides were eluted from the gel by incubating the sample overnight at 20°C under 600 rpm.

#### LC-MS/MS

1µg of the digested material was analysed using a UPLC nanoACQUITY (Waters, UK) coupled to a Q-Exactive Plus Hybrid Quadrupole-Orbitrap Mass Spectrometer (Thermo Scientific, USA). The chromatographic system is equipped with two columns. The first one, dedicated to the trapping of peptides, is a Symmetry C18 (5 µm, 180 µm x 20 mm, Waters, UK). The second one which performs the analytical separation is a HSST3 C18 (1.8 µm, 75 µm x 250 mm, Waters). The dimensions given for the columns are in the order: particle diameters, internal diameters and column lengths. Solvent A was water, acidified with 0.1% formic acid and solvent B was acetonitrile, also acidified with 0.1% formic acid. Cross-linked peptides were first trapped for 3min (98/2, v/v, A/B) at a flow rate of 20µL/min before being eluted with a gradient of 57 min at a flow rate of 700 nL/min. The elution started with a linear gradient of B from 2% to 7% in 5min, followed by an increase from 7% to 40% in 25min, then to 85% in 3min. This composition A/B 15/85 is kept for 5min before changing to 98/2 for reconditioning the analytical column. The Q-Exactive Plus spectrometer was set in a nanoESI positive mode acquisition for 57 minutes. The acquisition was recorded in full scan MS and data dependent MS/MS, in a mass range m/z 400-1,750. For the MS stage, resolving power was set at 70,000 @m/z 200, with an automatic gain control (AGC) target at 1e6 (or 50 ms as a maximum injection time). For MS/MS, a “Top 12” experiments was applied, meaning that the twelve most intense ions of each MS scan were selected for fragmentation. Singly charged ions, ions with undetermined charge (for example, electronic noise) and ions with signal intensities below the AGC threshold set at 1e3 were excluded from this selection. For precursor ions, the selection window was 2.0 m/z, the AGC target was 1e5 (or 50ms as a maximum injection time) and the resolving power of 17,500 @m/z 200. Normalized collision energy was 25. A dynamic exclusion of 10s was also applied to avoid the redundancy of MS/MS spectra of the same ions.

#### Data Analysis

Spectrum Identification Machine SIM-XL version 1.5.0.14^57^ was used for identification of cross-linked peptides. ADH-DMTMM crosslinker was set up in the software to create a mass shift of 138.0905 Da for each crosslink and 156.1012 for hydrolyzed monolinks. The possible reaction sites were restricted to C-terminal extremities and acidic side chains of Glu and Asp amino acids. Accuracy on mass measurements was 2 ppm for the precursor and 10 ppm for the fragment ions, to reduce to their maximum the false positives and to increase the reliability of the results. Oxidation of methionine, due to experimental conditions, has also been considered as a variable modification. Only the crosslinked peptides characterized by a SIM-XL internal score of 2.5, 3.0, 2.3, 3.0 or 2.5 and higher have been retained for IK, IJ, HJ, HI, or HK pairs, respectively. All the detected crosslinked peptides have been manually validated by analyzing their corresponding MS/MS spectra for exact masses, numbers of identified fragments, and signal intensity (*e*.*g*. **Figure 6b, Supplemental Figure 3b**).

### NMR experiments

All solution NMR experiments (except XcpH_p_/J_p_ chemical shift perturbation) were run at 37°C in a Bruker 900 MHz NMR spectrometer at the National Magnetic Resonance Facility at Madison. Sample condition used was NMR buffer containing 0.01% sodium azide, 50 µM 2,2-dimethyl-2-silapentane-5-sulfonate (DSS) (Sigma) and 8% D_2_O.

Backbone sequential assignments were carried out using standard 3D NMR experiments including HNCO, HN(CA)CO, HNCACB, CACB(CO)NH. These experiments were collected using non-uniform sampling (NUS). Data processing of NUS data was performed using NMRPipe^58^. Chemical shift referencing was done using DSS as reference. Backbone assignments were facilitated using NMRFAM-Sparky software^59^. Calculation of secondary chemical shifts for secondary structure estimation was done using random coil chemical shifts calculated for the XcpH amino acid sequence using ncIDP library^60^. Secondary chemical shifts were calculated as ΔδCa – ΔδCb, where ΔδCa = (δCa_XcpH_ – δCa_random coil_) and ΔδCb = (δCb_XcpH_ – δCb_random coil_). This step effectively removes any error in chemical shift referencing.

Chemical shift perturbation assays were carried out using 100 µM ^15^N-U XcpH and increasing amounts of unlabeled XcpJ (20 µM, 50 µM, and 80 µM). An independent ^1^H-^15^N HSQC spectrum was collected for each sample. Total change of the amide proton and nitrogen chemical shift (ΔδNH) was calculated for each residue as ((ΔδH)^2^ - 1/6 (ΔδN)^2^)^½^, were ΔδH and ΔδN correspond to the difference in chemical shift of proton and nitrogen respectively. ΔδNH was calculated using the control spectrum as the initial point and the spectrum with 80 µM XcpJ as the endpoint.

To perform paramagnetic relaxation enhancement (PRE) experiments, four different XcpJ_p_ cysteine mutant constructs (R46C, R53C, T178C, E180C) were used to attach the spin label (1-Oxyl-2,2,5,5-tetramethylpyrroline-3-methyl) methanethiosulfonate (MTSL) (Santa Cruz Biotechnology). XcpJ_p_ variants were purified as above in Buffer A containing 1 mM DTT. MTSL labeling of XcpJ_p_ was performed in NMR buffer and 0.5 mM DTT. XcpJ_p_ samples were incubated with a 20x molar excess of MTSL for 4 h at room temperature. Additional 20x molar excess of MTSL was added to the sample which was further incubated at room temperature overnight. Excess MTSL was removed by dialysis against the NMR buffer. The NMR sample contained 80 µM ^15^N-U XcpH_p_ and 24 µM of MTSL labeled XcpJ_p_. The ratio of XcpH to XcpJ_p_ was selected based on the chemical shift perturbation assay to avoid signal loss due to complex formation. A 2D ^1^H-^15^N HSQC spectrum was acquired for each sample. After collecting the HSQC experiment of the oxidized or paramagnetic form of MTSL, 2 mM sodium ascorbate (Sigma-Aldrich) was added to the sample and incubated at room temperature for at least one hour before collecting the spectrum of the reduced or diamagnetic MTSL form. Effects of the paramagnetic label were quantified as the ratio of signal intensity of the spectrum collected with MTSL in paramagnetic versus diamagnetic form (I_para_/I_dia_).

### Quaternary complex computational modeling

Modeling of the XcpH_p_I_p_J_p_K_p_ quaternary complex was achieved using high ambiguity driven protein-protein Docking (HADDOCK)^61^. Structures used for HADDOCK corresponded to the XcpI_p_J_p_K_p_ crystal structure (PDB: 5VTM) and an XcpH_p_ model prepared based on the structure of EpsH_p_ from *Vibrio cholerae* (PDB: 2QV8) using the Phyre2 server^62^. Allowed distances for acidic crosslinking restraints between C_*α*_ carbons were set between 8-18 Å, while 6-18 Å were used for PRE restraints (**Supplemental Tables 3 & 4**). Docking the XcpH_p_ model onto the XcpI_p_J_p_K_p_ structure was performed using the HADDOCK 2.2 server^61^. The best structure obtained from HADDOCK was selected for further analysis of the XcpH_p_I_p_J_p_K_p_ complex.

### Biological filament modeling

Modeling of the XcpG Type II secretion system pseudopilus was based on the PulG filament structure from *Klebsiella oxytoca*^15^ (PDB code 5WDA) using Pymol. Modeling was divided in the following steps: (1) adding missing transmembrane helices to XcpH_p_I_p_J_p_K_p_ proteins, (2) positioning the XcpHIJK complex onto the PulG filament (3) fitting XcpHIJK helices to the position of PulG transmembrane helices within the filament (4) fitting XcpG^63^ units to the PulG filament and (5) relaxing the whole system to remove atomic clashes. Missing transmembrane helices in the XcpH_p_I_p_J_p_K_p_ complex were initially modeled using the Phyre2 server using the respective amino acid sequence of the complete alpha helix. This modeled alpha helix was aligned to the soluble domain alpha helix structure in the HADDOCK model. Specific to XcpH, the helix was modeled on the PulG template from the EM reconstruction to include the unraveled section. Finally, the modeled helices were added to the XcpH_p_I_p_J_p_K_p_ complex PDB file using custom Python scripts. For XcpG a similar procedure was performed, in this case a model of the transmembrane helix was based on PulG helix (including the unraveled helix section), which was added to the XcpG_p_ solution NMR structure^63^ (PDB code 2KEP). The XcpHIJK complex with the helices added was positioned at the tip of PulG filament by aligning the XcpH soluble domain with the first PulG unit. Then, XcpHIJK model transmembrane helices were modeled semi-manually to fit the interior cavity of the PulG filament. Custom Python scripts were used to modify ϕ and ψ for this purpose. To guide the modeling, the transmembrane helices of PulG units were used as a reference. Finally, PulG subunits were replaced with XcpG to complete the pseudopilus filament. To achieve this, XcpG with the added helix was aligned to a PulG unit and XcpG helix was modeled to closely match the PulG transmembrane helix. Then, the modeled XcpG unit was copied and aligned to the PulG units to complete the filament. Finally, the whole model was relaxed with pyRosetta^52^. During this procedure, the known hydrogen bond between E5 carboxyl group and N-terminal amino group of the next subunit in the filament was enforced as a distance restraint of 2.8 Å. Other restraints to maintain the quaternary tip complex and XcpG filament assembly are shown **Supplemental Tables 5 & 6**. Ten filament models were produced, from which the best was selected based on favourable XcpG subunit packing and the lowest RMSD of the XcpH_p_I_p_J_p_K_p_ tip structure with respect the HADDOCK model.

## ACKNOWLEDGMENTS

NMR experimental data have been deposited to the BMRB with accession number 50449. We thank Dr. Eric Durand for pET-XcpG_p_ plasmid construction, Mrs. Nanou Tanteliarisoa for her help in the preparation of the samples dedicated to the acidic crosslinking, and Dr. Lisa Craig for valuable comments on the manuscript. We acknowledge funding from the ANR to RV (ANR-14-CE09-0027-01 & ANR-19-CE11-0020-01) and from the US National Science Foundation to KTF (IOS 1353674). Mass spectrometers used in this work were supported by the Walloon Region (Belgium) and the European Regional Development Fund. This study made use of the National Magnetic Resonance Facility at Madison, which is supported by NIH grant P41GM103399 (NIGMS), old number: P41RR002301. Equipment was purchased with funds from the University of Wisconsin-Madison, the NIH P41GM103399, S10RR02781, S10RR08438, S10RR023438, S10RR025062, S10RR029220), the NSF (DMB-8415048, OIA-9977486, BIR-9214394), and the USDA.

## AUTHOR CONTRIBUTIONS

RV, LQ, and KTF conceived the studies. CAE and BD subcloned, made site directed mutants, and purified soluble proteins. CAE carried out CSP and PRE experiments, and all quaternary structure and filament modeling. BD performed cysteine cross linking experiments. GB constructed *P. aeruginosa* strains and carried out secretion assays. SA carried out NMR experiments in Supp. Fig S4. BB was responsible for coevolution calculations in Supp. Fig S5. LQ performed all acidic cross linking methods development and experiments. RV, KTF, and CAE drafted the manuscript. All authors discussed and edited manuscript and prepared figures.

